# The frequency spectrum of chromatin accessibility in Yorubans points toward a significant role of random genetic drift in shaping the chromatin landscape

**DOI:** 10.1101/068718

**Authors:** Mariah Weavil-Abueg, Joshua G. Schraiber

## Abstract

The function of non-coding variation in the human genome is hotly debated. While much of the genome appears to be involved in some kind of molecular activity, a relatively small portion of the genome appears to be conserved across mammalian species. To try to understand part of this seeming paradox, we examined chromatin accessibility as a model molecular phenotype. We modeled chromatin state as either open or closed as looked at the frequency of open chromatin across 70 Yoruban cell lines. We saw that most regions of chromatin accessibility occurred in only a small number of individuals, although there are a number of regions that are accessible across the entire panel. To delve further into understanding the evolutionary mechanisms, we examined nucleotide diversity in and around accessible regions. We found that in the open chromatin access, low frequency regions had decreased nucleotide diversity, however, they were situated within regions of elevated nucleotide diversity. These results point toward a role of random mutation and genetic drift shaping the distribution of accessible regions in the human genome.

## Introduction

The vast majority of the human genome (∼95%) is made up of intergenic DNA. Several hypotheses exist to explain the preponderance of such non-coding DNA, ranging from neutral arguments that genome architecture is shaped by the mutation landscape [1] to functional hypotheses that this large amount of non-coding DNA allows for complex, modular and evolvable regulatory networks [2,3]. A key evolutionary prediction of the selective hypotheses is that intergenic DNA should show signatures of purifying selection, maintaining functional regions across evolutionary history; however, only approximately ∼5-10% of the genome seems to be evolving under purifying selection [4], which still leaves a large amount of non-coding DNA to be explained.

On the other hand, the functions of intergenic DNA may be highly buffered and turn over quickly, so looking directly at functional readouts is important for understanding the importance of non-coding DNA in humans. In 2012, the ENCODE consortium claimed that 80% of the human genome had some biochemical function [5]. They claimed that regions long thought to serve no biological role were important in shaping human phenotypes, which spurred controversy around what, exactly, is considered functional [6-8]. By definition, these regions do not code for proteins; however, they seem to be involved in some kind of transcriptional activity and/or DNA-protein interaction, suggesting a role in modulating gene expression.

One of the many factors affecting gene expression is the physical state of the DNA. While most of the genome is wound tightly around histone proteins (such regions are referred to as closed or inaccessible chromatin and likely to be transcriptionally silent, other areas are free of histones. Such regions of open, or accessible, chromatin, areas where the DNA is unwound, allow for proteins to bind to the DNA and the gene to be transcribed [9].

There is abundant evidence that chromatin accessibility both shapes and is shaped by transcriptional processes. For instance, transcription factors can prime accessible regions [10] and those open regions can direct the subsequent binding of downstream transcription factors [11].

Because chromatin accessibility varies across individuals in a population, there ought to be information about the causes and consequences of chromatin accessibility in polymorphism data. Recently, a group used DNase I hypersensitivity sequencing (DNase-seq) and mapped accessibility quantitative trait loci (dsQTL) [12]. Interestingly, they found that dsQTL significantly overlap with gene expression QTL (eQTL) of year-by genes, suggesting a role of chromatin variability among individuals in shaping expression variability among individuals. Moreover, frequently dsQTLs were seen within 200-300 base pairs of open chromatin regions, pointing toward the importance of cis-acting mutations in shaping both chromatin accessibility and gene expression.

Here, we revisit the data from Degner et al, identifying and characterizing differences in open chromatin regions in a population. By viewing chromatin accessibility as a high-dimensional phenotype, we can view each individual region of accessible chromatin as a correlated replicate of the same kind of evolutionary process, similar to how SNPs across the genome can be viewed as correlated replicates of the mutational and evolutionary process. This allows us to apply intuitions from population genetics directly to the phenotypes themselves, which may yield insights that would be difficult to obtain by examining sequence variation alone.

## Materials and Methods

### DNase data

We downloaded publically accessible DNase I hypersensitivity data from 70 Yoruban lymphoblastoid cell lines, which has been aligned by Degner et al. Mapped reads were stored in a custom format by Degner et al, and were converted to .bed files using a custom script.

### Calling DNase I Hypersensitivity Peaks

On the basis of a recent analysis [13] we used Hotspot [11] to detect regions of open chromatin. We used an FDR of 0.01 and used the mappability files for 22bp reads provided by the authors of Hotspot. We merged peaks across individuals using bedops [14].

### Peak Trimming

Merging peaks across individuals can result in artificial overlap in DHS across individuals by inadvertently merging long stretches of nearby peaks. To compensate, we imposed two filters on all of our peaks. First, we required that merged peaks don’t merge two peaks that were called separately in a single individual. Second, we examined the length of the merged peaks. Figure 1A shows that empirical cumulative distribution function of merged peak lengths, while Figure 1B shows how many individuals are repeated in a given peak as a function of its length. On the basis of examining these figures, we chose the require peaks be less than 500bp. In total, we had 1,176,504 DHS across the 70 individuals, which was reduced to 772,617 peaks with a mean length of 270.3bp after trimming.

**Figure 1.**
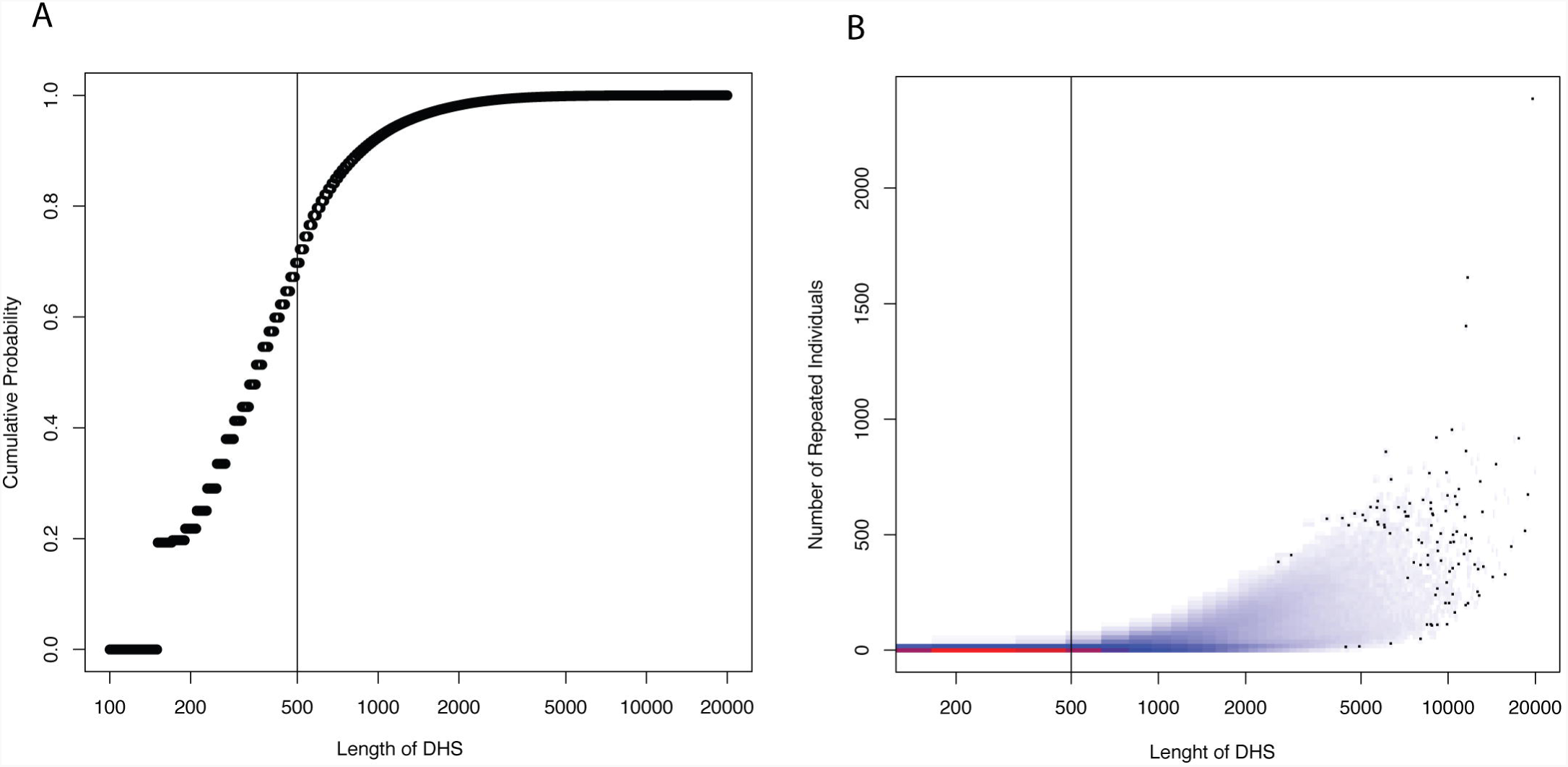
Panel A: Empirical cumulative distribution function of lengths of merged peaks. The x axis shows peak length, while the y axis shows the proportion of peaks with that length or less. Solid vertical line indicates the cutoff of 500bp that we chose for filtering peaks. Panel B: Smoothed scatter plot of number of repeated individuals per peak against merged peak length. Red indicates high density, blue indicates low density. An individual is counted as repeated if at least two distinct peaks from the single-individual peak calling show up as part of one peak in the merged call.

### Population genetic analysis

To examine population genetic properties of chromatin accessibility polymorphism, we computed the nucleotide diversity for each polymorphic DHS we identified. We downloaded the 1000 genomes phase 3 data [15], and restricted our analysis to Yoruban individuals. We then computed nucleotide diversity for a given DHS as

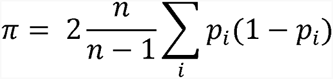

where *i* ranges over all variable sites in the DHS, n is the number of individuals, and *p_i_* is the frequency of polymorphic site *i*. We computed nucleotide diversity in two types of windows: in one situation, we restricted the calculation to within the merged DHS, and in the other we expanded 200bp outside of the called peak.

## Results

We first computed the frequency distribution of regions of accessible chromatin that passed our filters (Figure 2). The frequency spectrum mostly displays a negative slope, with higher frequency DHS occurring more and more rarely. However, there is an enrichment of regions that are found to be open in all individuals (lower right corner).

**Figure 2.**
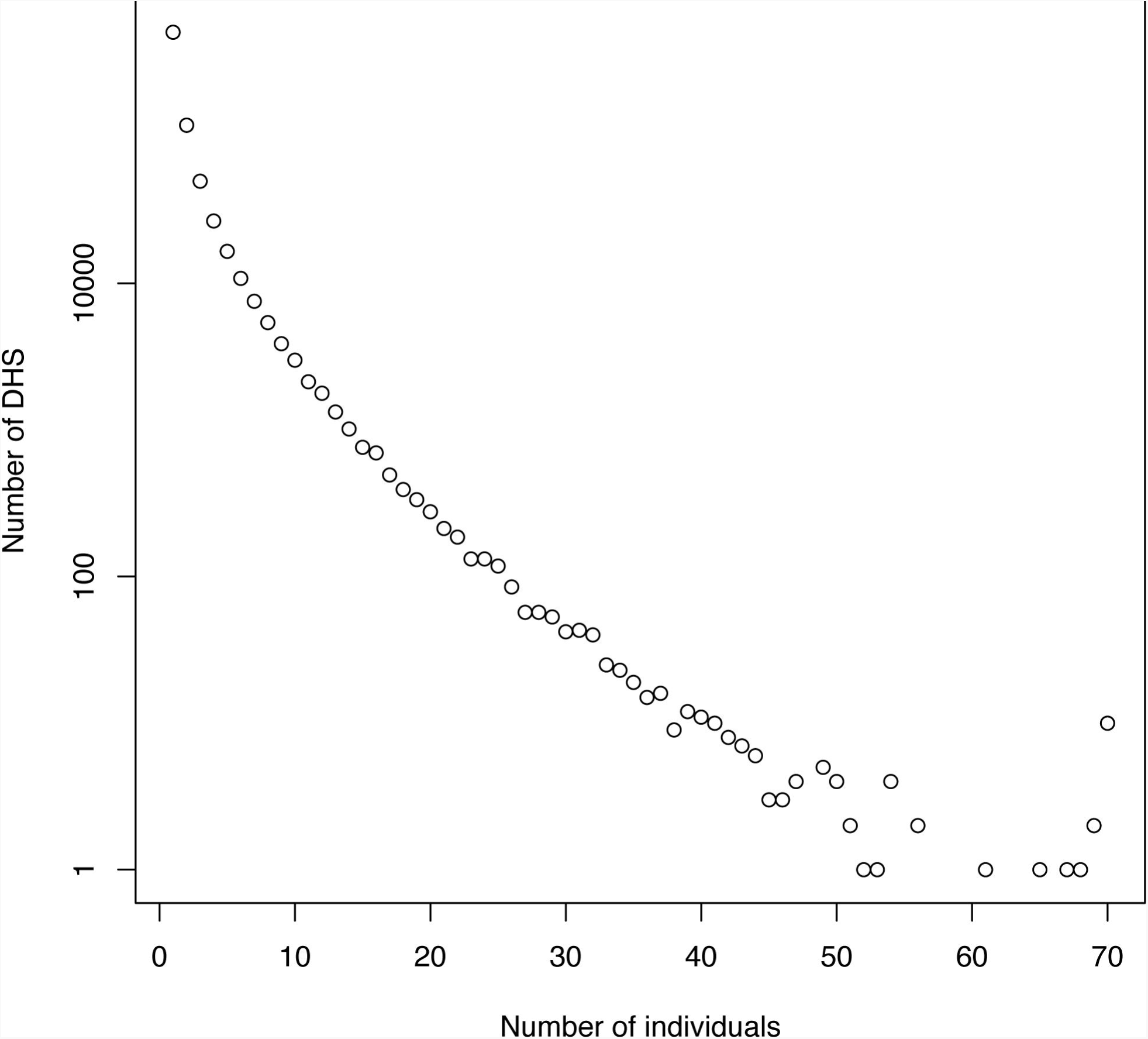
Frequency spectrum of accessible chromatin. The x axis is the number of individuals in which a region is called accessible, and the y axis is the number of regions with that frequency of accessibility. Note that y axis is log scaled.

We next examined the population genetic properties of regions of accessible chromatin that occur at different frequencies. We found contrasting patterns depending on whether we examined polymorphism exclusively within the DHS or including 200 base pairs of surrounding sequencing. When we look at just the merged peak region, we see that there is lower nucleotide diversity in regions of lower frequency DHS (Figure 3A). However, when including 200bp of sequence context, we see higher nucleotide diversity in regions where the fewest individuals share a region of accessible chromatin (Figure 3B). In both cases, there appears to be a trend toward lower nucleotide diversity as the frequency of open chromatin reaches middle frequencies, although there are so few high frequency regions that there is not much signal for them.

**Figure 3.**
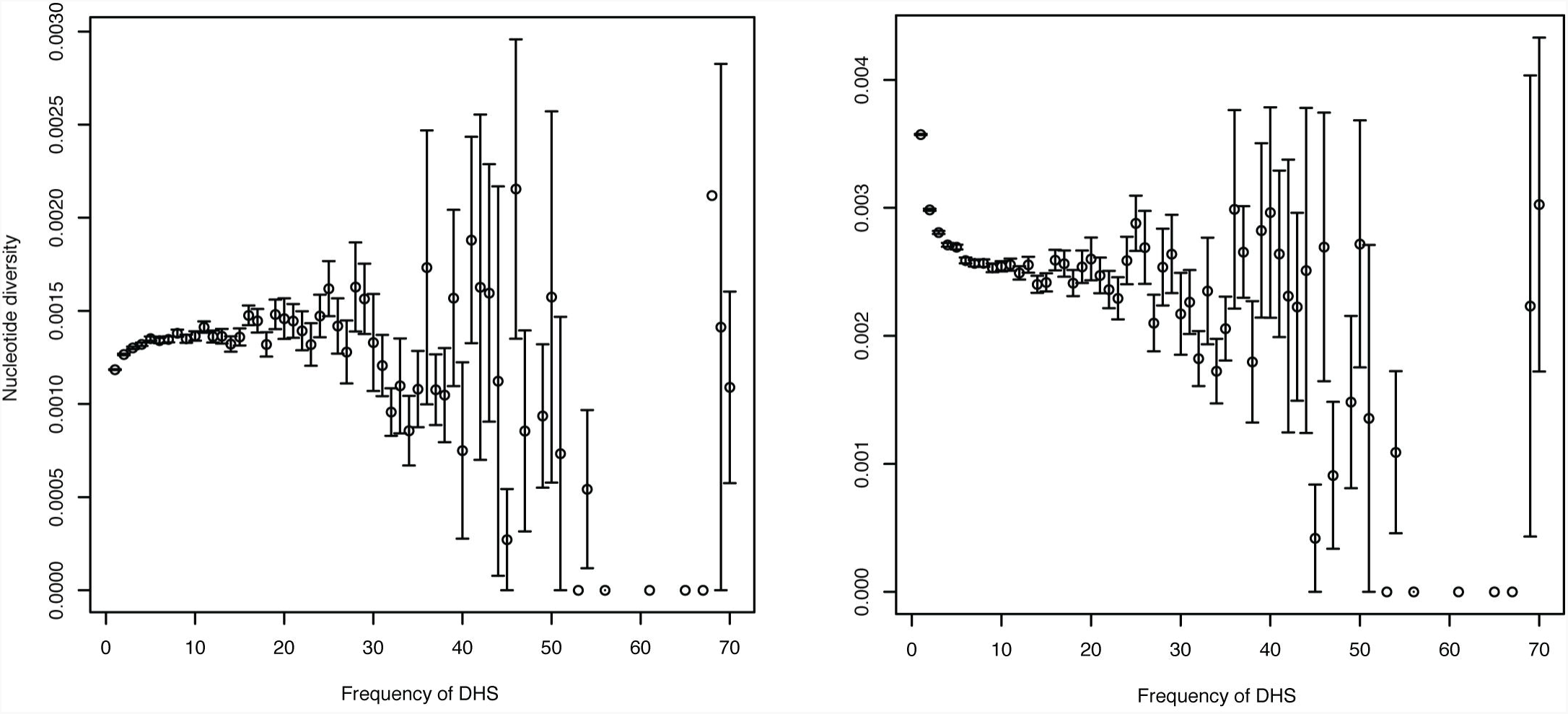
Nucleotide diversity in and around accessible regions of different frequencies. In each panel, the x axis shows the frequency of the accessible region and the y axis shows the nucleotide diversity. Panel A shows nucleotide diversity estimated only within the peak region, while Panel B shows nucleotide diversity estimated in the peak region and 200bp of context on either side.

## Discussion

Despite making up a vast majority of the human genome, the importance of non-coding DNA is poorly understood. Although much of the genome is involved in some kind of measurable molecular activity [5], the extent to which that activity represents an important function is unknown.

One important window into the fitness consequences of non-coding variation is examination of nucleotide polymorphism that may underlie molecular phenotypes. To that end, several studies have identified QTL for various aspects of molecular activity, including gene expression, chromatin accessibility, and DNA-protein interactions [12,16-20]. By analyzing variation in and around QTL for molecular phenotypes, inferences can be made about the evolutionary importance of these loci, and hence the importance of the traits they control.

The high dimensional nature of molecular activity also makes studying phenotypes in-and-of-themselves a valuable tool for understanding their fitness consequences. This kind of approach has been successful in analyzing gene expression evolution in molecular pathways [21-24] adapted this idea to think of each individual region of accessible chromatin as a two-state discrete phenotype and examined polymorphism in chromatin accessibility across 70 Yoruban cell lines.

Interestingly, we found that most DNase-I hypersensitivity sites occur at a low frequency in these cell lines. This is suggestive that many regions of accessible chromatin may be simply a result of random effects, either due to low frequency mutations changing DNA-histone interactions or simply rare, stochastic biophysical effects. We do see a spike of DHS that occur in every individual assayed, which is suggestive that those may be critical to cellular function.

To gain some insight into the functional importance of regions of accessible chromatin that occur at different frequencies across individuals, we computed nucleotide diversity across the different frequency categories. Unexpectedly, we found that whether we focus explicitly on the DHS region or include 200 base pairs of context made a strong difference in findings. When considering the region of open chromatin itself, we found that low frequency regions had decreased nucleotide diversity. This observation ran contrary to our expectation that such low frequency regions may be due to random mutations in more polymorphic regions of the genome. However, when considering 200 base pairs of context, we found that lower frequency regions of open chromatin indeed had elevated levels of nucleotide diversity. This may suggest a complicated role of mutational context in shaping the landscape of low frequency chromatin accessibility.

As we expected, regions with a higher frequency of chromatin accessibility seem to show a trend for lower levels of nucleotide diversity, suggesting that they are more highly constrained. However, the pattern is difficult to discern due to the low number of high frequency accessible regions. More sensitive measurements of chromatin accessibility may lead to overall more regions detected and may increase the power to understand the mechanisms underlying the maintenance of high frequency DHS.

Our results suggest a key role for random processes in shaping the chromatin landscape in humans. This does not necessarily imply that chromatin accessibility is by-and-large a neutral trait. In particular, there may be modules of coordinated regions of chromatin, of which only a subset of regions are required to be open in order for downstream functions to occur. In this case, any individual region may appear to evolve neutrally, when not considered in the context of the larger module. Future work should consider patterns of chromatin accessibility in physically proximal regions, as well as incorporate information about known enhancers and their downstream targets.

Although we motivated our work by appealing to the standard models of population genetics, we did not actually pursue any theoretical models. A promising direction for the future is to attempt to model chromatin accessibility variability in a population from first principles, for example by using a threshold model [25].

By integrating insights from molecular and evolutionary studies, we can make progress toward an understanding of non-coding variation in humans as well as further details about the molecular features that lead to human specific traits.

## Acknowledgements

We thank Joshua Akey for helpful discussions about this work.

